# Population specific reference panels are crucial for the genetic analyses of Native Hawai’ians: an example of the *CREBRF* locus

**DOI:** 10.1101/789073

**Authors:** Meng Lin, Christian Caberto, Peggy Wan, Yuqing Li, Annette Lum-Jones, Maarit Tiirikainen, Loreall Pooler, Brooke Nakamura, Xin Sheng, Jacqueline Porcel, Unhee Lim, Veronica Wendy Setiawan, Loïc Le Marchand, Lynne R. Wilkens, Christopher A. Haiman, Iona Cheng, Charleston W. K. Chiang

## Abstract

Statistical imputation applied to genome-wide array data is the most cost-effective approach to complete the catalog of genetic variation in a study population. However, imputed genotypes in underrepresented populations incur greater inaccuracies due to ascertainment bias and a lack of representation among reference individuals,, further contributing to the obstacles to study these populations. Here we examined the consequences due to the lack of representation by genotyping a functionally important, Polynesian-specific variant, rs373863828, in the *CREBRF* gene, in a large number of self-reported Native Hawai’ians (N=3,693) from the Multiethnic Cohort. We found the derived allele of rs373863828 was significantly associated with several adiposity traits with large effects (*e.g.* 0.214 s.d., or approximately 1.28 kg/m^2^, per allele, in BMI as the most significant; P = 7.5×10^−5^). Due to the current absence of Polynesian representation in publicly accessible reference sequences, rs373863828 or any of its proxies could not be tested through imputation using these existing resources. Moreover, the association signals at this Polynesian-specific variant could not be captured by alternative approaches, such as admixture mapping. In contrast, highly accurate imputation can be achieved even if a small number (<200) of Polynesian reference individuals were available. By constructing an internal set of Polynesian reference individuals, we were able to increase sample size for analysis up to 3,936 individuals, and improved the statistical evidence of association (e.g. p = 1.5×10^−7^, 3×10^−6^, and 1.4×10^−4^ for BMI, hip circumference, and T2D, respectively). Taken together, our results suggest the alarming possibility that lack of representation in reference panels would inhibit discovery of functionally important, population-specific loci such as *CREBRF*. Yet, they could be easily detected and prioritized with improved representation of diverse populations in sequencing studies.

## Introduction

Statistical imputation of untyped variants is a crucial step for large-scale genetic investigations of complex traits. By comparing to an appropriate reference panel, often based on whole genome sequences of individuals, statistical imputation infers the genotype at variant sites not covered on genotyping arrays^1^.Therefore, imputation benefits many study cohorts with its balance between budget and coverage of the genome. Nevertheless, inadequacy and inaccuracy of imputed markers can impede downstream genetic studies or clinical screening. This problem could arise when reference panels are not genetically close to the population of interest, further exacerbated by ascertainment bias of existing genotyping array. Today, it is increasingly recognized that there has been a severe bias towards studying individuals of European origin in GWAS ^2–5^, and similarly in the largest available reference panel for imputation (*e.g.* the Haplotype Reference Consortium^6^). The inability to statistically impute diverse populations further hinders progress in studying these diverse populations that otherwise are already underserved ^7,8^.

One of the major obstacles in studying diverse, non-European, populations, particularly for indigenous communities such as the Native Hawai’ians, is the inability to accrue large sample sizes^9^. For example, GWAS in European ancestry-based cohorts numbers in greater than 1 million^10^; By contrast, there are only ∼1.2 million individuals in total living in the United States that may derive some part of their ancestry to Native Hawai’ians, according to the US 2010 census survey (https://www.census.gov/prod/cen2010/briefs/c2010br-02.pdf). Therefore, the focus in studying diverse populations is often in (1) evaluating the transferability of findings from large-scale European ancestry-based studies, and (2) identifying population-specific variants that may contribute to genetic risks in non-European populations. As they are often very rare or missing in Europeans, population-specific variants in non-Europeans are usually absent in most genotyping arrays. These variants would rely on high quality imputation in order to be captured, unless the population of interest is whole genome sequenced at large scale. One recent example of a variant that bears important consequences to the health and disease risk of a population is the Polynesian-specific missense variant rs373863828 in gene *CREBRF*. This locus was initially detected because of an association signal of a proxy variant, rs12513649, which was on the Affymetrix 6.0 array. The missense rs373863828 was then discovered through targeted resequencing of a small number of private Samoan sequences, followed by imputation into the entire Samoan cohort and validation of the imputed genotypes. Rs373863828 was found to have a large effect on body mass index (BMI) as well as on a number of other adiposity, metabolic, and anthropometric traits in Samoans ^11^. Despite an estimated 26% allele frequency in Samoans, the derived allele is only segregating in the few Pacific Islands populations and not found elsewhere in the world ^12–16^. Because of a lack in Polynesian haplotypes in publicly accessible sequencing databases (e.g. 1000 Genomes Project or Haplotype Reference Consortium), this variant could not be directly imputed and studied by researchers with publicly available resources.

In this study, we use the *CREBRF* locus as an example to examine the potential limitations of post-imputation analyses in the absence of a proper representation in reference panels. We genotyped the variant in Native Hawai’ians from the Multiethnic Cohort (MEC). The Hawai’i archipelago in North-East Polynesia was first settled between 1,200 ∼1,800 *ya* ^17–20^. Historically, Native Hawai’ians have remained relatively isolated on the northern Pacific islands, until their recent encounter with inter-continental migrants from Europe around the late 18^th^ century, and from East Asia (mainly China and Japan) during the 19^th^ −20^th^ century, followed by minor contributions from other populations around the world^17,18^. Compared to other populations, contemporary Native Hawai’ians have higher incidence rate of obesity-related medical conditions, such as diabetes and cardiovascular diseases^21^. Therefore, it is of clinical importance to characterize the impact of variants with potentially large effects on adiposity, such as rs373863828 in *CREBRF*, in this population. We demonstrate that despite a strong impact on adiposity, the *CREBRF* locus could not be discovered using conventional mapping methods with currently available resources, predicting important challenges for discovering additional variants contributing to population-specific genetic risks. However, our findings also suggest that these challenges could be mitigated if the representation in reference sequences were improved, even quite marginally.

## Methods

### Study subjects

The Multiethnic Cohort (MEC) is a population-based cohort study that examines lifestyle and genetic risk factors for cancer. It consists of 215,251 adult men and women from Hawai’i and California (primarily Los Angeles County), with ages ranging from 45 to 75 at recruitment (1993-1996). The cohort includes mainly five ethnicities: Native Hawai’ians, African Americans, Japanese Americans, Latinos, and European Americans. Participants entered the cohort by completing a questionnaire on diet, height, weight, demographic information, and other risk factors. Participants are followed up with questionnaires every five years, and linked with cancer registries annually. More details of the MEC can be found in Kolonel et al. ^22^ The institutional review boards of the University of Hawai’i and the University of Southern California approved the study protocol. All participants signed an informed consent form.

### Genotyping and QC

In total, 4,990 MEC Native Hawai’ian participants were genotyped by genome-wide SNP arrays across different arrays platforms. Among them, 3,940 individuals were genotyped on Illumina MEGA array as part of the PAGE consortium^3^; 307 were genotyped on Illumina MEGA^EX^ as part of a collection of additional obesity related anthropometric and imaging measurements of adiposity^23^; An additional 266, 318, and 492 individuals were genotyped on Illumina Infinium Oncoarray, Illumina Human 1M Duo BeadChip, and Illumina 660W arrays, respectively, in the past for studies of colorectal cancer, nicotine metabolism, and breast cancer^24–26^. In the main text and Table S1, we refer to these past studies using MEC Native Hawai’ian samples as PAGE, Obesity, CRC, smokers, and NHBC substudies, respectively. In this report, we focused most of the analyses on Native Hawai’ian participants in the PAGE consortium as this study had the largest sample size and SNP density, compared to other studies.

The genotyping calling process and quality control filtering for all of the genotyped datasets above are described in the corresponding references, with exception of the obesity related study on MEGA^EX^. In this dataset, DNA extraction from blood samples for the MEC-Adiposity Phenotype Study participants^23^ was performed using the Qiagen QIAmp DNA kit (Qiagen Inc., Valencia, CA). DNA samples were genotyped by the Illumina MEGA^EX^ array. SNPs with a call rate <0.95, a replicate concordance <100% based on 39 QC replicate samples, and those with poor clustering after visual inspection were removed. Problematic samples with a call rate <0.95 or gender/sex mismatches were removed.

We applied additional uniform quality controls as follows: All variant names were updated to dbSNP v144; duplicated loci and indels were removed; triallelic variants or variants with non-matching alleles to 1000 Genomes Project phase 3 (1KGP)^27^ were discarded; loci with unique positions not found in 1KGP were removed from the dataset; alleles were standardized to the positive strand by comparing to 1KGP. Finally, a genotype missingness filter of 5% was applied.

Additionally, we genotyped rs373863828 in *CREBRF* in a total of 4,331 MEC Native Hawai’ians using a Taqman Assay; genotypes for 4,214 of these individuals were called successfully, 3,693 of which also have genome-wide array data and thus formed the dataset of this study. This variant was also genotyped in an additional 407 European Americans, 313 African Americans, 432 Japanese Americans, and 386 Latinos (both American and non-American born Hispanics) from the MEC; the variant was monomorphic in all these other populations examined.

### Phenotypes analyzed

We focused on a total of 30 quantitative and dichotomous traits related to obesity, type-2 diabetes, and cardiovascular diseases, chosen because Native Hawai’ians have shown to have excess risk for these traits compared to other populations^21,28,29^. These include 13 quantitative traits (BMI, hip circumference, waist circumference, adult standing height, waist-hip ratio, total cholesterol, fasting glucose, adiponectin, HDL, LDL, hypertension, fasting insulin and HOMA-IR) and 2 dichtomous traits (obesity, type-2 diabetes). We also analyzed an additional 10 adiposity traits measured using whole-body dual-energy X-ray absorptiometry (DXA) and abdominal magnetic resonance imaging (MRI) for a subset of 307 individuals^23^: total fat mass, total lean mass, lean mass in leg, lean mass in arm, percentage of total fat, trunk fat mass, percentage of liver fat, visceral fat area, and abdominal fat area. Finally, we examined 5 additional dichotomous disease traits related to cardiovascular outcomes in MEC participants enrolled in the Medicare fee-for-service program including: heart failure (HF), hyperlipidemia (HYPERL), hypertension (HYPERT), ischemic heart disease (IHD), and stroke/transient ischemic attack (TIA). These phenotypes were identified by CMS Chronic Conditions Data Warehouse using algorithms that search the Medicare claims data for specific diagnosis or procedure codes (https://www2.ccwdata.org/web/guest/condition-categories). Please refer to Table S2 for details of the phenotype transformation.

### Imputation of rs373863828

In order to impute rs373863828 using a population-specific reference panel, we identify MEC Native Hawai’ian individuals with the highest amount of Polynesian ancestry to serve as the Polynesian reference panel. Specifically, we first estimated the global ancestry proportions of 3,940 subjects from PAGE, using ADMIXTURE v1.3^30^. We modeled the Native Hawai’ian ancestry as 4 ancestral components: the majority Polynesian component, with recent admixtures from Europeans, East Asians, and Africans. We thus included 1000 Genomes Project ^27^(1KGP; Phase 3) GBR, CEU, TSI, IBS as European references; CHB, JPT, CHS, CDX, KHV as East Asian references; YRI and LWK as African references. After pruning the dataset to exclude loci with genotype missingness >5% and minor allele frequency <1%, we stratified individuals into a group of closely related 1^st^ or 2^nd^ degree relatives estimated from KING^31^ and a separate group of relatively unrelated individuals. We then pruned variants such that no two variants have an LD above r^2^ of 0.1, per recommendation of ADMIXTURE, and conducted an unsupervised run across the unrelated individuals at *k*=4. We then projected the estimated ancestral allele frequency to the related samples to infer the genomic ancestries of these individuals. We performed five independent iterations with randomized seed numbers, and found no minor mode at *k*=4. The final result was integrated by averaging the estimated proportions after matching ancestry clusters across five runs. From this analysis we identified 178 MEC Native Hawai’ian individuals with >90% Polynesian ancestry and have a refined kinship coefficient (through PC-relate^32^, below) < 0.2 among reference samples. We further subset to 152 individuals who were genotyped successfully at rs373863828. To construct the imputation reference panel, the genotype of the rs373863828 was merged to the array genotypes on chromosome 5 for these 152 individuals and merged with the 1KGP, assuming that each 1KGP sample carries homozygous reference genotype at rs373863828. We statistically phased this constructed reference panel again with EAGLE2, as rephasing was shown to improve imputation accuracy^6^ and we found that rephasing improved genetic ancestry inference (Figure S9). We used minimac3^33^ (version 2.0.1) and the constructed reference panel to impute the genotype of rs373863828 in Native Hawai’ian individuals with genome-wide array data. After imputation, we had genotype dosages of rs373863828 available for all samples, covering those phenotyped but not particularly genotyped by Taqman assay, which increased the sample size in the refined associations of this locus with all traits available.

### Constructing the genetic relatedness matrix

Due to sample relatedness and population substructure within the MEC Native Hawai’ians, standard approaches for constructing principal components and kinship estimates could each bias one another. Thus, we used PC-air and PC-relate from GENESIS v2.4.0 package^32,34^, which performs a principal component analysis robust to family structures, and infers genetic relatedness unbiased from unspecified population structures, respectively. Based on our initial kinship estimates from KING^31^, we obtained the top 10 eigenvectors reflecting ancestry influences at the default unrelatedness cut-off of 2^(−11/2). The unbiased eigenvectors were in turn used to refine the kinship coefficients in the genetic relatedness matrix (GRM).

### Linear mixed model

We used a linear or logistic mixed model (LMM) implemented in EMMAX^35^ to perform all association tests in this study. We used the GRM generated from PC-relate (above) as a random effect in the model, and the inverse normalized residuals and covariates as fixed effects for each trait (Table S2). In association tests of the *CREBRF* region on chromosome 5, only 15,334 markers that were genotyped or had imputation INFO score > 0.4 were included. In admixture mapping with BMI and T2D, genotyped positions with probabilities of being on Native Hawai’ian haplotypes, as inferred from local ancestry inference (below), were used as dosages. Bonferroni correction was directly applied to associations conducted in PAGE subjects. For the additional 10 adiposity traits in the separate 307 individuals, due to the likely correlations among the traits, we determined the number of independent tests as following: we decomposed the phenotypic matrix by principle component analysis, and calculated the accumulative variance explained by eigenvectors until it surpass 95% of that of the phenotypic matrix. We found the corresponding number of PCs surpassed the threshold to be 7, i.e. the number of independent tests considered for the associations with the 10 adiposity traits.

### Local ancestry inference

We used RFMix2^36^ to estimate local ancestry on the rephased genotype data of MEC Native Hawai’ians from the PAGE dataset. We used our constructed imputation reference panel as reference panel of the four components of ancestry. HapMap2^37^ pooled recombination rate (ftp://ftp-trace.ncbi.nih.gov/1000genomes/ftp/technical/working/20110106_recombination_hotspots/) was used as the genetic map. We adopted the default parameters of RFMix2 as we found no notable difference when enabling expectation-maximization for 5 iterations, or when enabling re-analysis of reference individuals (data not shown). From the output of RFMix tsv file we also computed global ancestry estimates for each person after we excluded tracts with any ancestry probability lower than 0.9. The global ancestry proportions estimated from RFMix is highly concordant with those from ADMIXTURE (Figure S9); the main deviation is due to individuals detectably related to the 178 reference individuals (maximal kinship coefficients with the 178 internal individuals are significantly higher than the rest; Mann-Whitney (one way) P =5.14E-58).

### Power estimate of single variant association with BMI

We estimated the power of discovering a locus rs373863828 via single variant association, using BMI as an example. We followed a standard power estimate for quantitative traits: assuming a chi-square model with degree of freedom as 1 applies, the power equals to the left tail probability of a chi-square value of corresponding alpha probability, but with the non-centrality parameter shifted as the product of sample size and heritability explained by the single locus. For rs373863828,

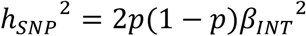

Where p is the MAF (6%), and *β*_*INT*_ is the effect size in standard deviation unit (0.248).

### Power simulation for admixture mapping

We assumed the following in our power simulation: 1) the derived allele frequency of rs373863828 in the ancestral, unadmixed, Native Hawai’ians is 13%, as estimated from current individuals who have >90% Polynesian ancestry; 2) the effect size of the derived allele and the phenotypic standard error are transformed to the same units as reported in the discovery cohort in Samoans by Minster et al.^11^, i.e. 1.36 and 6.9 kg/m^2^ ; 3) the percentage of local ancestry at *CREBRF* region among all samples is similar to the average of global ancestry proportions (*i.e.* no strong selection at the locus) (Figure S4).

Given a target sample size, we assigned the genotype of each individual assuming Hardy-Weinberg equilibrium and the derived allele frequency of 5.9% (matching the frequency for rs373863828 in Native Hawai’ians). Given the genotype, we then assigned the ancestral origin of haplotype for each individual. If an individual has:

1. Homozygous derived genotype implies the individual derived both alleles from Polynesian haplotypes;
2. Heterozygous genotype implies the individual carries at least one copy of Polynesian haplotype. For the other, non-derived, allele, the probability that this allele derives from a Polynesian haplotype is:

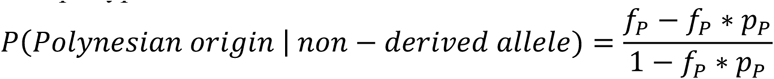

where *f*_*P*_is the global Polynesian ancestry percentage, and *p*_*P*_is the derived allele frequency in non-admixed ancestral Polynesians (13%). Thus, probability of this individual carries exactly one copy of Polynesian haplotype is 1 − *P(Polynesian origin* | *non* − *derived allele*).
3. Homozygous ancestral genotype (GG): the number of Polynesian tracts each individual carry corresponds to a binomial sampling at the probability of P(Polynesian origin | non-derived allele) as described, and the number of trials is 2 (diploid).

To simulate phenotypes of individuals given their genotypes, we sampled from a normal distribution with mean shifted by 2β, β, and 0 standard deviations, where β is the reported effect size. Given simulated genotype, local ancestry, and phenotype, we tested the association between local ancestry and the phenotype to simulate admixture mapping. Power was calculated as the number of times a positive association at or below the specified statistical threshold was achieved in 500 iterations.

### Selection test

We performed nS_L_ scan^38^ implemented in Selscan^39^, which identifies ongoing positive selection in genome based on phased haplotypes and is robust to recombination rate variation, on the 152 internal reference individuals at the *CREBRF* locus. To standardize the nSL score at the *CREBRF* locus, we constructed the null distribution based on genome-wide nSL scores from variants with derived allele frequency between 12-14%.

## Results

### The missense variant in CREBRF is correlated with the proportion of Polynesian ancestry

To characterize the functional missense variant, rs373863828-A, in Native Hawai’ians, we genotyped this single locus in 3,693 self-reported Native Hawai’ian individuals from the Multiethnic Cohort (MEC), in addition to their existing genome-wide array data (Table S1). We also genotyped this variant in 1,538 individuals from other continental populations also found in the MEC (Methods). Consistent with previous reports that this variant is found exclusively in Pacific Islanders^11,12^, we estimated the derived allele frequency to be 5.9% in Native Hawai’ians, but is monomorphic in all other ethnicities we genotyped. As Native Hawai’ians derive a large proportion of their ancestry from Polynesians, we also found significant correlations between the derived allele frequency (DAF) at rs373863828 and individuals’ indigenous Polynesian ancestry proportions (r = 0.98, p = 6.2e-7, Figure S1; ancestry binned in 10 percentage points), and between the direct genotypes and individual ancestry proportions (GLM β=2.67, p < 2e-16). Individuals who are relatively unadmixed (with estimated Polynesian ancestry >90%) carry the allele at the frequency of 12.8% (Figure S1).

The lower frequency (12.8%) among Native Hawai’ian individuals who are relatively unadmixed is unexpected, given that the allele has been reported to be under positive selection and is segregating at 26% in Samoans^11^. We attempted to replicate the signal of selection at this locus in the Native Hawai’ians using the nSL (ref ^38^) among the 152 individuals with estimated Polynesian ancestry > 90%. Compared to randomly drawn variants throughout the genome matched by derived allele frequency, we found no evidence of rs373863828-A being positively selected (nSL = 0.72, p = 0.57, Figure S2).

### The CREBRF variant is associated with adiposity traits in Native Hawai’ians

The *CREBRF* variant has been reported to have a large effect on body mass index (BMI) in several populations from the Pacific Islands, and significantly associated with height and other adiposity traits in Samoans^11,40^. To explore its impact in Native Hawai’ians, where the derived allele is present at a relatively lower frequency compared to the Samoan discovery cohorts (26%), we conducted linear mixed association tests for the variant among genotyped individuals who also had available a collection of quantitative anthropometric, metabolic, and lipid phenotypes (Table S2 and Table S3). After phenotype transformations (Table S2), we replicated the increasing effect of the derived allele for rs373863828 on BMI (β=0.214 s.d. per allele, P = 7.55e-5) and height (β=0.182 s.d. per allele, P = 3.96E-4). Based on these genotyped Native Hawai’ians, this variant explains 0.52% and 0.37% of the phenotypic variance in BMI and height, respectively. These are much larger than the largest effect loci found in Europeans for these traits (in Europeans, rs11642015 in *FTO* and rs143384 in *GDF5* explain 0.25% and 0.2% of the variation for BMI and height, respectively, based on ∼360,000 individuals studied in UK BioBank; http://www.nealelab.is/uk-biobank/). We also found the derived allele was associated with increases in waist and hip circumferences (β=0.215 s.d. per allele and 0.206 s.d. per allele, P = 8.7E-4 and 0.00127, respectively), but was not associated with waist-hip ratio (Table S4). Consistent with the Samoan study, we found no association between rs373863828-A and several metabolic and serum lipid traits, including fasting insulin, HOMA-IR, adiponectin, HDL, LDL, or triglycerides (Figure 1, Table S4). In addition, we did not find an association of rs373863828 with total cholesterol or HDL, which was reported otherwise in Samoans^11^.

**Figure 1.**
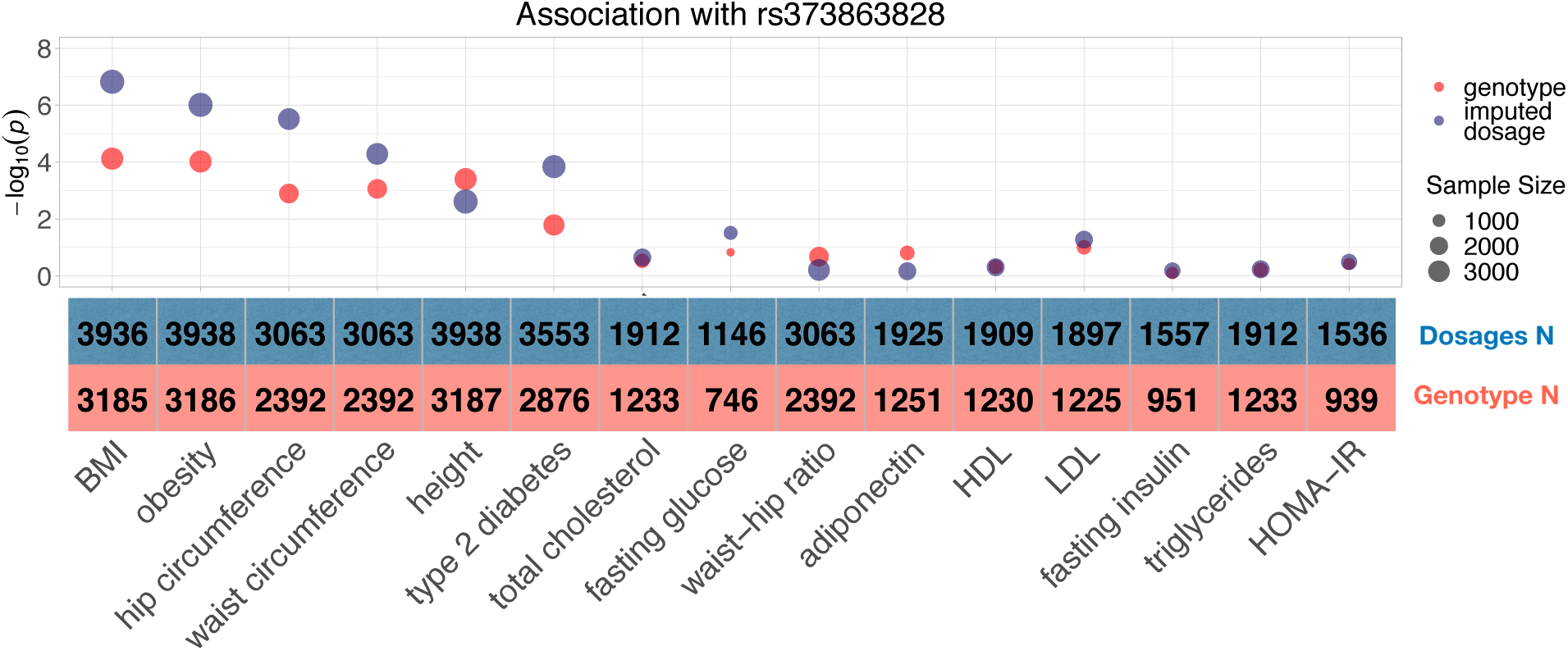
Association of rs373863828 with adiposity and lipid traits from PAGE cohort in Native Hawai’ians. Log-transformed P values from association of rs373863828 with adiposity- and lipid related traits. Associations with direct genotypes and imputed dosages are denoted in red and blue, respectively. Point size is proportional to the sample size in each test, and also shown below the trait.

We also examined the association for a number of dichotomous disease phenotypes. As expected for its large effect on BMI, we found the derived allele to increase the risk for obesity (OR=1.096, P = 9.52E-5). In addition, we replicated the variant’s protective effect to risk of type-2 diabetes (T2D) (OR=0.935, P = 0.0162). As obesity is a major risk factor for cardiovascular diseases and the Native Hawai’ians exhibit excess risk for cardiovascular diseases when compared to Europeans^28,29^, we also tested the effect of the derived allele of rs373863828 on five traits among the Medicare FFS participants for the same cohort: heart failure (HF), hyperlipidemia, hypertension, ischemic heart disease (IHD), stroke/transient ischemic attack (TIA), but we found no significant associations regardless of whether or not we controlled for BMI in the analysis (Table S5).

Finally, to examine the effect of this allele on more refined measures of body fat distribution, we tested the association of the derived allele with ten additional adiposity traits collected on a subset of 307 Native Hawai’ian individuals in our study^23^. While the variant was significantly associated with overall fat mass and whole body fat percentage (β=0.69 s.d. per allele and 0.58 s.d. per allele, P = 0.001 and 0.007, respectively, after multi-trait test correction), the role it has on fat distribution is less clear to characterize, as the signals were nominally significant, or marginal on nominal significance with lean mass in arms and legs (P = 0.056 and 0.013, respectively), and subcutaneous fat mass (P=0.088), but not associated with other adiposity traits (Table 1).

**Table 1.** Associations of rs373863828 with adiposity traits measured from DXA and abdominal MRI.

| Trait | Typed genotype |  |  | Imputed dosages |  |  |
| --- | --- | --- | --- | --- | --- | --- |
|  | Sample size | Effect size<sup>1</sup> | P value | Sample size | Effect size<sup>1</sup> | P value |
| Total fat mass | 294 | 0.69 | 0.001 | 298 | 0.68 | 0.001 |
| Fat percentage | 294 | 0.575 | 0.007 | 298 | 0.565 | 0.008 |
| Trunk fat percentage | 291 | 0.085 | 0.672 | 295 | 0.08 | 0.691 |
| Abdominal fat | 291 | 0.13 | 0.303 | 295 | 0.131 | 0.301 |
| Liver fat percentage | 276 | 0.148 | 0.497 | 280 | 0.159 | 0.465 |
| Subcutaneous fat | 278 | 0.349 | 0.088 | 282 | 0.335 | 0.1 |
| Visceral fat | 278 | 0.118 | 0.571 | 282 | 0.12 | 0.564 |
| Lean mass in arm | 283 | -0.399 | 0.056 | 287 | -0.396 | 0.057 |
| Lean mass in leg | 288 | -0.507 | 0.013 | 292 | -0.507 | 0.012 |
| Total lean mass | 292 | 0.233 | 0.233 | 296 | 0.229 | 0.242 |

<sup>1</sup>Effect size is based on inverse normalized transformation of original phenotypes (Methods), and are thus in units of standard deviations.

### GWAS with current imputation resources or admixture mapping are unlikely to discover the CREBRF variant

While we have generally replicated the large effect rs373863828 exerts on BMI and other adiposity traits in the Native Hawai’ians, approaches to discover variants like this also exemplifies one of the main goals in genetic studies of diverse populations. The derived allele of rs373863828 has large effects and is population specific, suggesting that genotype and/or ancestry at this locus is important for risk assessment in the Native Hawai’ian and other Pacific Islanders. Using *CREBRF* as a test case, we thus examined whether a similar locus like this could have been discovered using currently available resources. If the entire cohort were whole-genome sequenced, we estimated moderate power to identify this variant: at genome-wide significance threshold (i.e. 5E-8), we have 41% power with the current sample size (N=3940). The power at the same significance threshold is much greater (75%) with the entire MEC Native Hawai’ian group (N∼ 5400 individuals), a sample size comparable to the Samoan discovery population sample^11^ that first reported this variant.

However, it is not yet feasible to sequence the whole-genome of all MEC Native Hawai’ians. Therefore, statistical imputation is the most efficient strategy for gene discovery. Currently, 1000 Genomes Project (Phase 3; 1KGP) is the most diverse public sequencing database. Because rs373863828 is absent in 1KGP, it cannot be imputed directly. Yet there is the possibility to impute a proxy variant nearby that could tag the causal variant. We thus imputed the array genotype data in all Native Hawai’ian samples using 1KGP data (Figure S8), and conducted a scan for association across the *CREBRF* locus using the linear mixed model for BMI or type II diabetes as examples of quantitative or dichotomous trait. However, we found no hint of association around the *CREBRF* region for either phenotype (+/- 100kb; lowest *P* = 1.2E-4 and 2.5E-3 for BMI and T2D, respectively, compared to a significance threshold of 5e-8) (Figure S3), suggesting that the current imputation resource is not sufficient for detecting this locus in Native Hawai’ians.

We then examined the imputation quality of a known proxy variant rs12513649, with which the initial association in Samoans led the researchers to hone in on resequencing the entire *CREBRF* locus^11^. Rs12513649 is in high LD with rs373863828 (r^2^ >= 0.988) in the Samoans, and similarly in 178 unadmixed Native Hawai’ians (r^2^ >0.99)(Methods). As this variant segregates in 1000 Genomes East Asians at ∼ 6%, in theory it is reasonably imputable. However, the overall imputation quality at rs12513649 is below standard pre-GWAS QC (MiniMac R^2^ <0.4). Moreover, despite 80% of imputed genotypes at the locus have posterior genotype probability (GP) >0.9, this observation of high GP was driven by the homozygous ancestral genotype; the confidence of imputed genotype dropped sharply among carriers of derived allele at either the proxy (rs12513649) or the missense (rs373863828) variant (14.0% and 52.8%, respectively, Table S6).

An alternative approach to discover this locus in the Native Hawai’ians would be to take advantage of the recent admixture and conduct admixture mapping. Given locally resolved assignment of ancestry across the genome in an admixed population, admixture mapping tests the association of local ancestry with a quantitative or dichotomous phenotype. There is reasonable *a priori* expectation that admixture mapping could successfully identify the *CREBRF* locus for its association with BMI or T2D, given that it is population specific, correlates strongly with Polynesian ancestry (Figure S1), and exerts a large effect on BMI, which is differentially distributed between ancestral populations. However, we found no significant association via linear mixed models with Polynesian ancestry across the gene (P = 0.057 for BMI, 0.452 for T2D, compared to a conventional statistical threshold of 5e-5 for admixture mapping). We estimated the discovery power for this locus, given the current sample size of 3,940 and other assumptions of the allelic effect (Methods), to be as low as 0.2% for a P <= 5E-5 threshold (Figure 2, Figure S5). In fact, even at 5% type I rate of replication standard, the power of replicating the *CREBRF* region via admixture mapping is only 18.2% (Figure S6).

**Figure 2.**
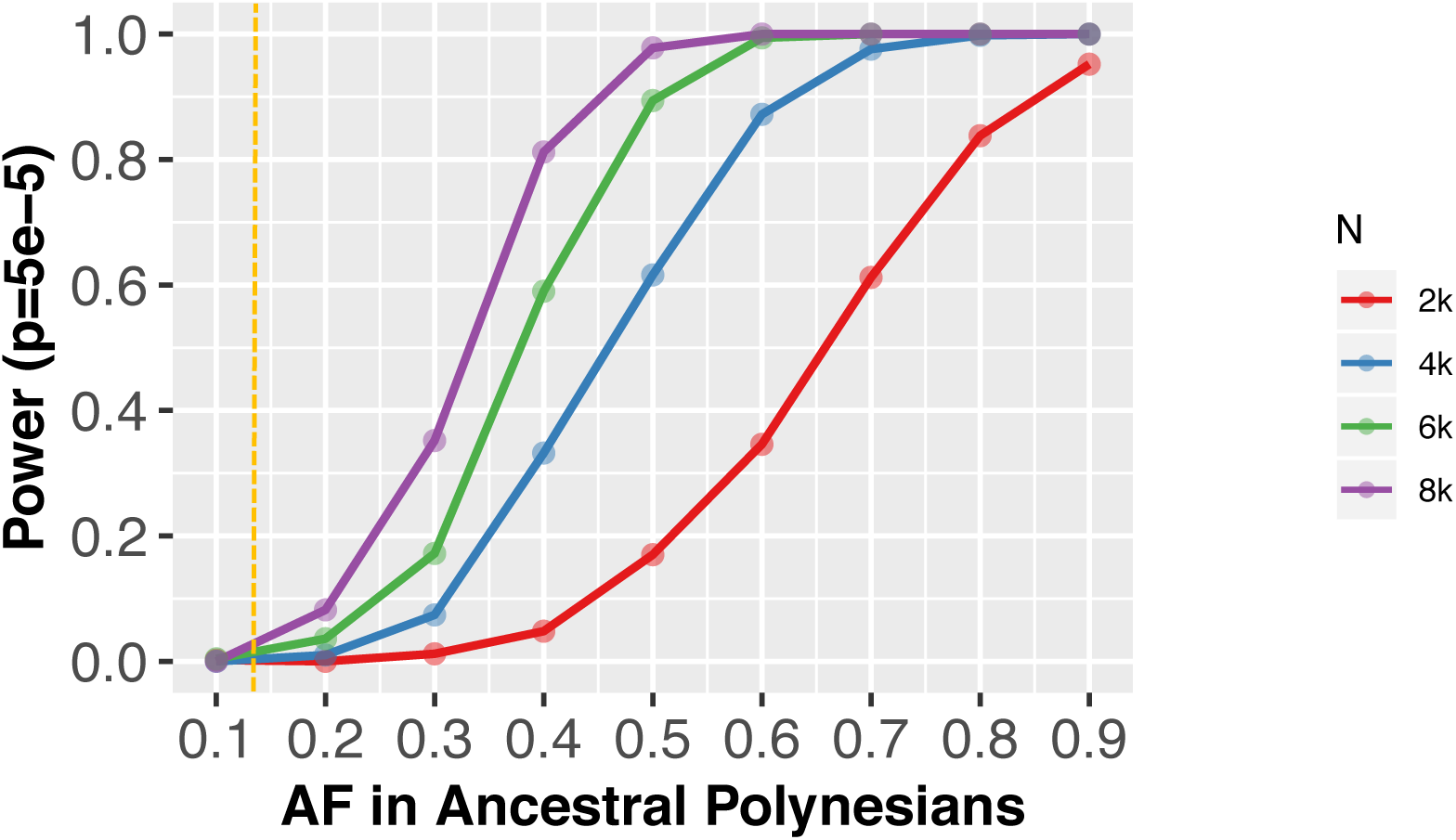
Statistical power of admixture mapping to discover *CREBRF* with BMI. Power was estimated through simulation given a range of allele frequencies in ancestral Polynesians and sample sizes. The effect sizes were assumed using rs373863828 as example, and the yellow dashed line denotes the empirical allele frequency of rs373863828 in unadmixed Native Hawai’ians (Methods). The significance threshold for genome-wide discovery via admixture mapping is set at 5e-5.

Taken together, without an appropriate imputation reference panel for Native Hawai’ians, viable alternative approaches currently available could not have efficiently mapped this locus.

### Imputation against internally constructed Polynesian reference boosts association signals

The obstacles described so far to conduct genetic analysis in diverse population are much attributed to the lack of representation from diverse populations in public whole genome sequences. We thus simulated a situation where a small number of individuals were available as a part of the reference panel to test if the key variant, rs373863828, could be imputed well and mapped in association studies (Figure S7). We selected 152 unrelated, relatively unadmixed (> 90% indigenous ancestry) individuals who had rs373863828 successfully genotyped to be our Polynesian reference, and merged the genotype data with 1KGP. Using this reference panel, we imputed the genotype of rs373863828 among all samples on the 5 different array platforms (Figure 3, Table S7). Regardless of the platform a dataset was genotyped on, we found the imputed dosages highly correlated with the directly assayed genotypes at rs373863828: r^2^ > 0.92 in all datasets tested, except for one dataset genotyped on the Oncoarray (0.76; Table S7). Other measurements of imputation quality, including concordance rate and allele frequency based internal R^2^ from Minimac^33^, also supported the high imputation accuracy (Table S7). While we included all relatively unadmixed individuals as the reference panel, we also found that imputation quality remained largely unchanged (Table S9) when a set of randomly selected individuals of the same size were used as reference panel, despite the allele segregating at lower frequency (Figure 1).

**Figure 3.**
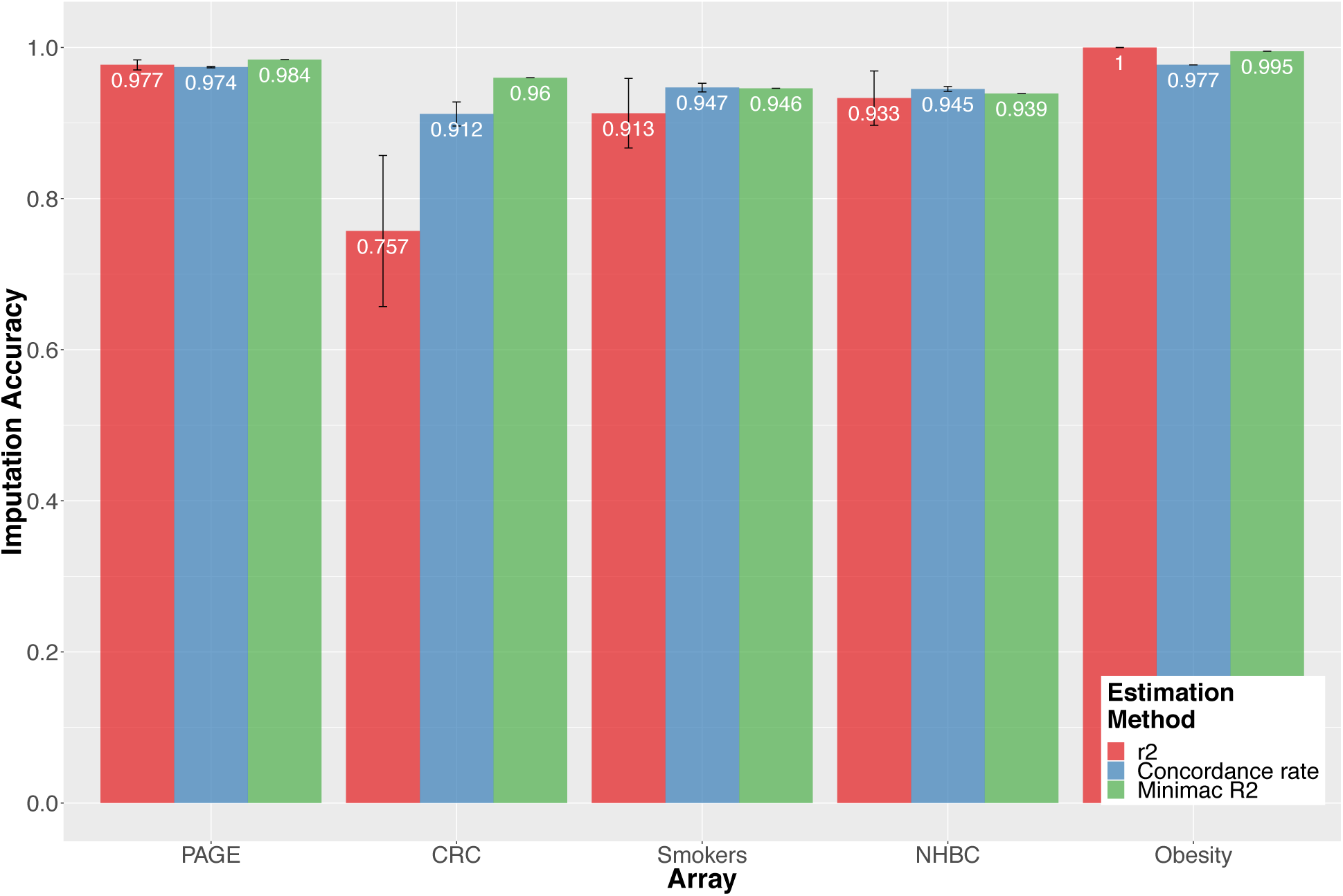
Assessment of imputation accuracy of rs373863828 using internally constructed references. The reference panel included 152 unadmixed Native Hawai’ians, and other populations from 1000 Genome Project (Methods). Imputation accuracy was estimated by comparing the true genotypes at rs373863828 and the imputed dosages, via different estimation methods. Standard errors were obtained through bootstrap.

Refining the associations with traits using imputed dosages of rs373863828, we found the statistical evidence for association in traits that were previously significant or marginally significant to generally improve: A larger sample size was achieved to include all phenotyped subjects, as compared to only a subset of individuals were particularly genotyped on the *CREBRF* missense variant in each study (Table S1). These were reflected mostly in adiposity phenotypes, such as BMI (P = 1.5E-7), waist and hip circumference (P = 5.15E-7 and 3.07E-6, respectively), and in dichotomous traits, obesity (P = 9.7E-7) and type II diabetes (P = 1.43e-4). Fasting glucose, previously non-significant in the genotype association, would now surpass nominal significance level after imputation (P = 0.031, Figure 1, Table S4).

## Discussion

We demonstrated the urgent need of having proper reference sequences in order to explore population-specific variants in diverse populations by using rs373863828 in *CREBRF* as an example in this study. We replicated the increasing effect of the derived allele of this variant on anthropometric and adiposity traits in Native Hawai’ians, and its protective effect on type II diabetes, consistent with reports in other populations from the Pacific Islands^11,15,16^. When examining more refined measure of body fat distribution, we also found the derived allele to be associated with increasing total fat mass and whole body fat percentage, even though we only have data available on ∼300 Native Hawai’ians. Most importantly, using the *CREBRF* locus in Native Hawai’ian as an example, we have shown that even though this locus is exhibiting some of the largest effects on BMI observed in humans, its poor coverage in publicly available reference database due to the population specificity prevents the efficient mapping of this locus. Alternative mapping strategies such as admixture mapping also would not be powered enough to identify this locus. Then, without a specific staged study design to investigate diverse populations, we would not have been able to identify this variant that might contribute to health disparity between populations.

While our findings largely support those reported in the Samoans^11,40^, we did not replicate the reported association with total cholesterol, even after imputing the variant in all individuals with phenotype available. For fasting glucose, the association was also only nominally significant after imputation. This may suggest that the allelic effect is potentially mediated by environmental factors that are found in the Samoans only, or potentially more likely, the sample sizes with these traits available in the Native Hawai’ians are still insufficient.

We were also unable to detect the reported signature of natural selection at the *CREBRF* locus in Native Hawai’ians. Consistent with this observation is the lowered derived allele frequency in Native Hawai’ians (approximately 13% in relatively unadmixed Native Hawai’ians, vs. 26% in Samoans). While the settlement in Polynesia is believed to have occurred in a west-to-east direction across the Pacific, the derived allele is also found in lower frequency (∼2 to 19%) in other populations in Pacific, including Tongans and New Zealand Maori living west of Samoa^12–15^. It is unclear whether the difference of allele frequencies among the Pacific Islanders is mostly attributed to the differences in selection strength, the bottleneck and genetic drift in the founding Polynesians, different admixture histories, or some combinations of the above. A more detailed explanation will require a better construction of demographic history of the different groups.

The opposite effect on obesity and type II diabetes, two typically comorbid conditions, suggests that rs373863828-A could play a role in fat distribution among different body areas. One possible explanation is that the derived allele promotes accumulation of subcutaneous fat mass better than that of visceral fat, as the former can lead to obese phenotype, while excess of the latter, other than general adiposity, is associated with insulin resistance and contributes to peripheral insulin sensitivity^41,42^. To test this hypothesis, we examined the association with subcutaneous and visceral fat measures available in 278 Native Hawai’ians, controlling for their total fat mass. We were not powered to identify significant association with either trait, although the estimated effect size was larger for subcutaneous fat than for visceral fat. Further investigations are needed to explore the role of rs373863828 in general on fat deposition to better understand obesity-related metabolic disease.

Finally, of most immediate concerns is the need to construct population specific reference panels to aid further genetic analysis of these populations. Again taking the rs373863828 as an example, the variant is not found in either the 1000 Genomes Project or the Haplotype Reference Consortium, because the variant is exceedingly rare outside of Polynesia and some other Pacific Islands (gnomAD frequency of 3e-5), and neither of the reference datasets contained Polynesians. The recent release of the Human Genome Diversity Project^43,44^ contained Pacific Islanders, but the variant was also not found probably due to small sample sizes (N=28). A subset of the Samoan cohort was sequenced as part of the TOPMed consortium^45^; therefore, public release of this reference data for imputation will help if we can assume population continuity between Samoans, Native Hawai’ians, and their most recent common ancestors. Alternatively, another approach to overcome the issue of under-representation for investigators is to sequence a small number of individuals within the population of interest; we have shown that a sample size less than 200 is likely to be adequate for accurate imputation of this locus. Sequencing data, complemented by efforts to expand the cohort, would have the potential to detect other population-specific alleles of importance to health disparity.

In summary, there is an urgent need to increase diversity in public genome sequence reference panels. The current deficiency could lead to adverse consequences such as failing to discover population-specific risk variants. This is particularly important for rare variants with potentially large effects, as rare variants tend to be geographically restricted and yet bear a significant role in understanding genetic architectures amongst populations^3,46,47^. As sequencing cost continue to decline, it will be more feasible to establish diverse, population-specific references, empowering investigations for these functionally important variants like rs373863828 in the *CREBRF* gene.

## Author contributions

C.W.K.C., I.C., C.A.H., L.L.M. provided project supervision; C.Caberto, A.LJ., M.T., L.P., B.N., G.S. performed genotyping experiments and quality control; C.Caberto, P.W., J.P., V.W.S., Y.L., U.L., L.R.W. performed phenotype data harmonization; M.L. conducted analyses; M.L., C.W.K.C. wrote the manuscript with input from all co-authors.

## Acknowledgements

We would like to thank all MEC Native Hawai’ian participants involved in this study. We acknowledge Ryan L. Minster at University of Pittsburgh and Stephen T McGarvey at Brown University for their helpful advice on the manuscript. This study was funded by U01 CA164973, P01CA168530, U01HG007397, and UH Cancer Center Support Grant for Genomics and Bioinformatics Shared Resource P30CA071789.

## Supplementary Data

Figure S1. Derived allele frequency at rs373863828 binned by Polynesian ancestry proportions. Figure S2. nSL summary statistics

Figure S3. Locuszoom of *CREBRF* region 1KGP3-imputed dosages and (A) BMI. (B) T2D Figure S4. Local ancestry plot at CREBRF and Chromosome 5

Figure S5. Power simulation under a range of Polynesian ancestry (discovery level)

Figure S6. Power simulation under a range of Polynesian ancestry (replication level)

Figure S7. Imputation schema with internally constructed panel

Figure S8. Imputation schema with 1KGP3.

Figure S9. Global ancestry comparison between ADMIXTURE and RFMix

Table S1. Sample size and genotyping information across cohorts

Table S2. Phenotype transformation and covariates for linear mixed model associations.

Table S3. Summary of untransformed quantitative phenotypes used in association analyses

Table S4. Associations of rs373863828 with adiposity, metabolic, and serum lipid traits in PAGE cohort

Table S5. Associations of rs373863828 with cardiovascular diseases in Medicare FFS Table S6. Imputation quality of rs15213649 across array platforms

Table S7. Imputation R^2^ at rs373863828 against internal reference

Table S8. Imputation R^2^ at rs373863828 against random individuals as reference

---

**Figure S1.**
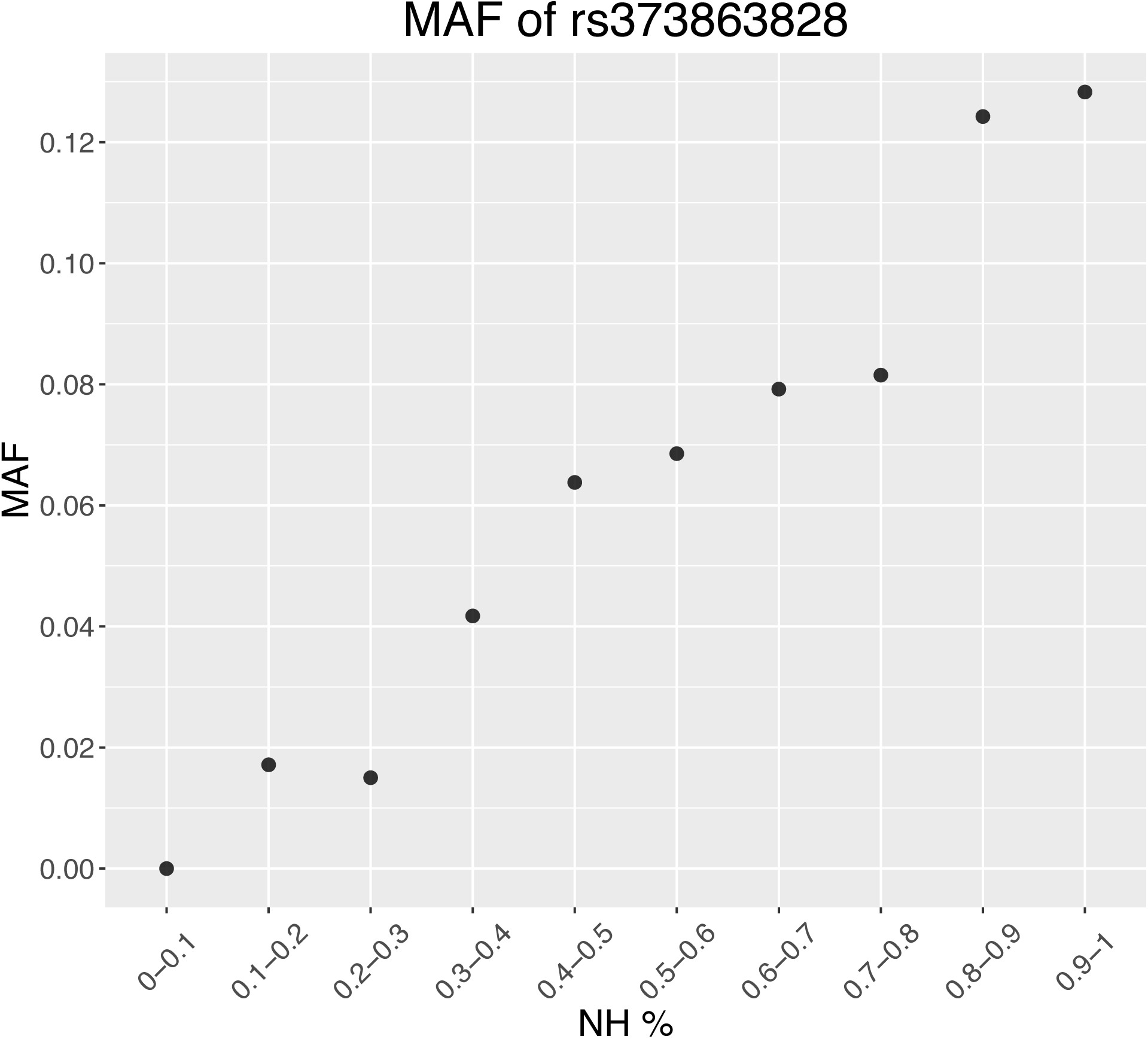
Derived allele frequency at rs373863828 calculated in all individuals genotyped at the locus, binned by Polynesian ancestry proportions estimated by ADMIXTURE.

**Figure S2.**
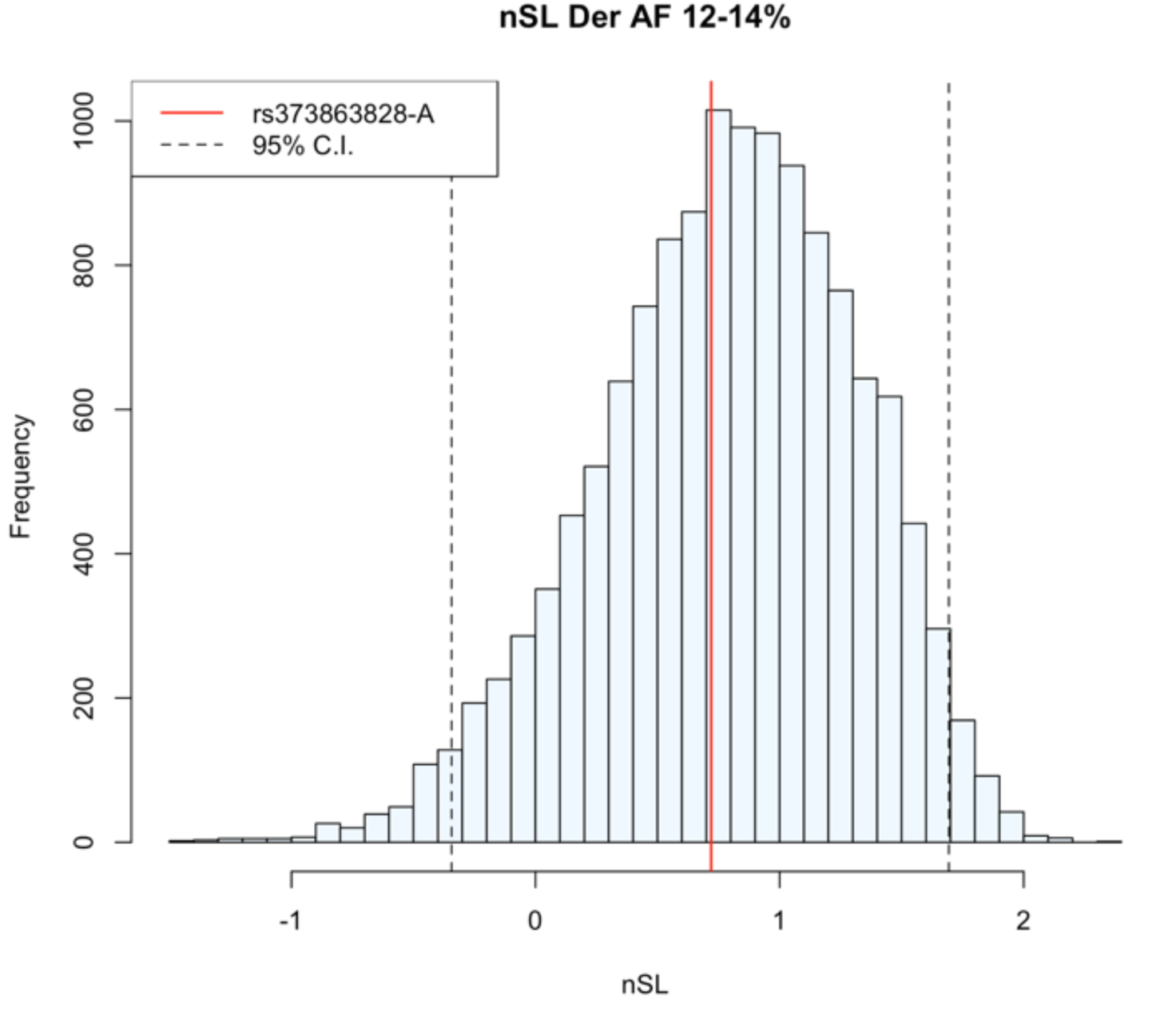
nSL summary statistics in 178 unrelated Native Hawaiians with indigenous ancestry >90%. The null distribution (blue) was formed by nSL calculated on genome-wide derived alleles whose frequencies are between 12%-14%. Red line denotes nSL of rs373863828 (MAF=13%). Dashed lines represent the 95% confidence intervals.

**Figure S3.**
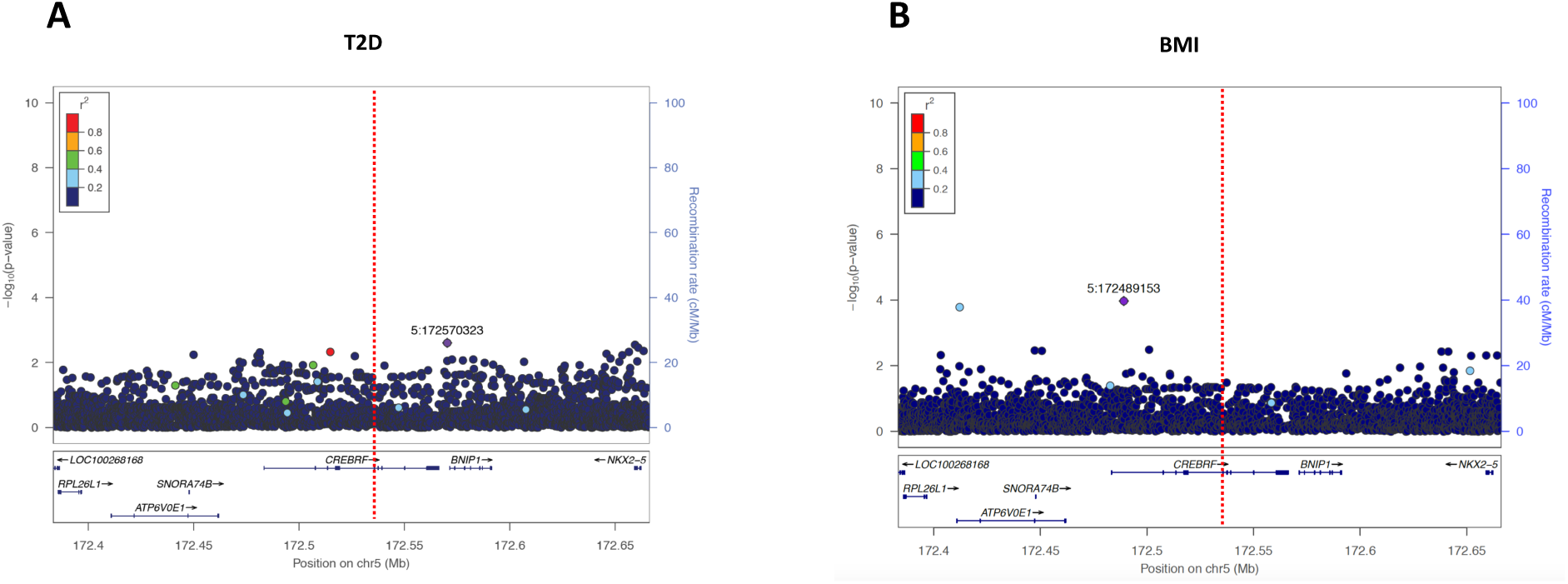
Locuszoom of *CREBRF* region (+/- 100 kb) for association with (A) BMI and (B) T2D. Imputed dosage using 1KGP as the reference panel was used in association testing. The variants with the lowest P value are denoted with their rsID, Red dashed line represents the position of rs373863828.

**Figure S4.**
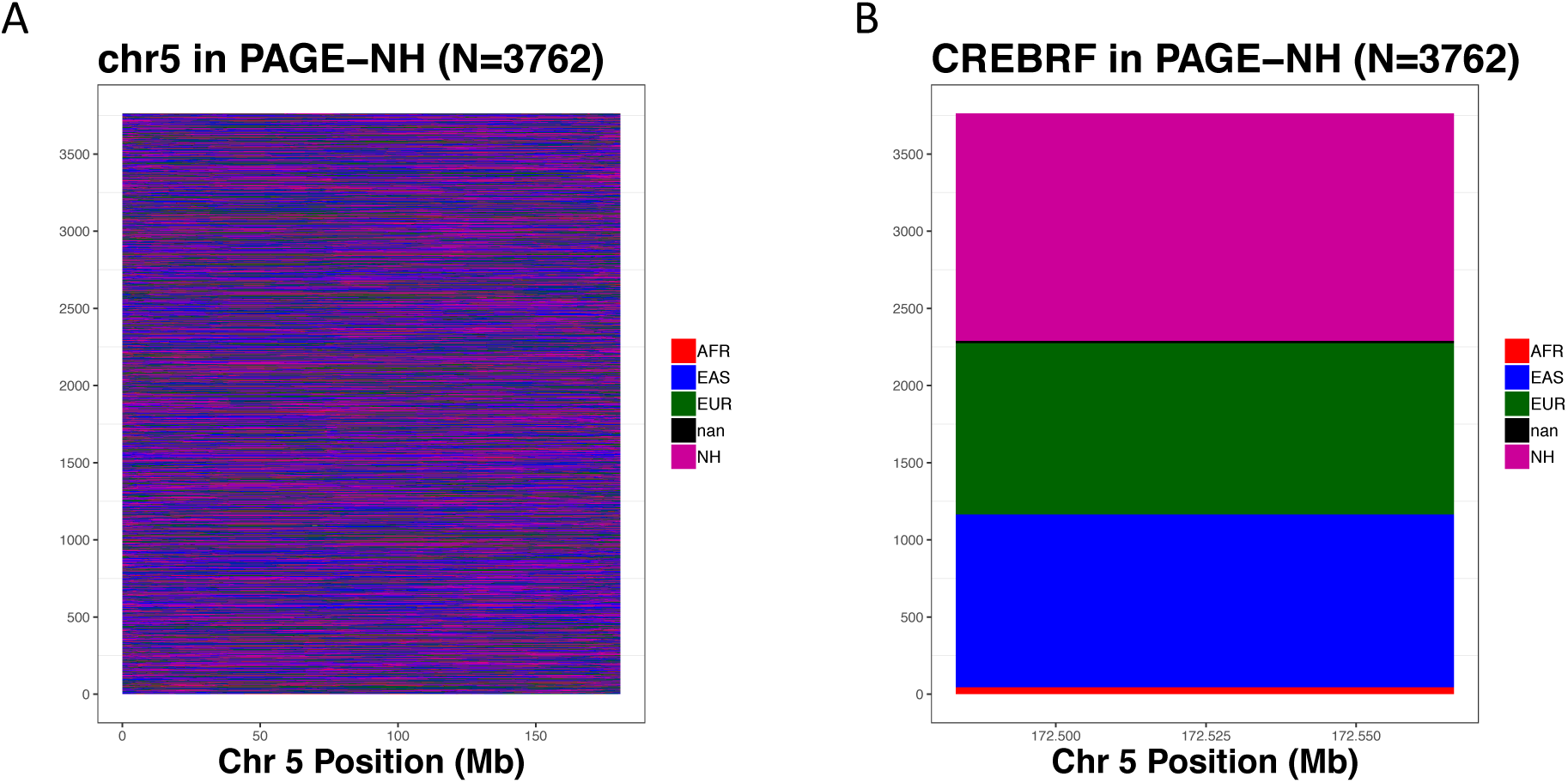
Local ancestry inference of (A) the entire Chromosome 5 and (B) the *CREBRF* region, excluding the 178 individuals with Polynesian ancestry > 90% that was used as reference. Each row corresponds to a single phased haplotype, and is sorted in (B) by ancestry. Each line of haplotype is colored based on its inferred ancestry, where black (“nan”) represents no call, with maximum inferred probability < 0.9.

**Figure S5.**
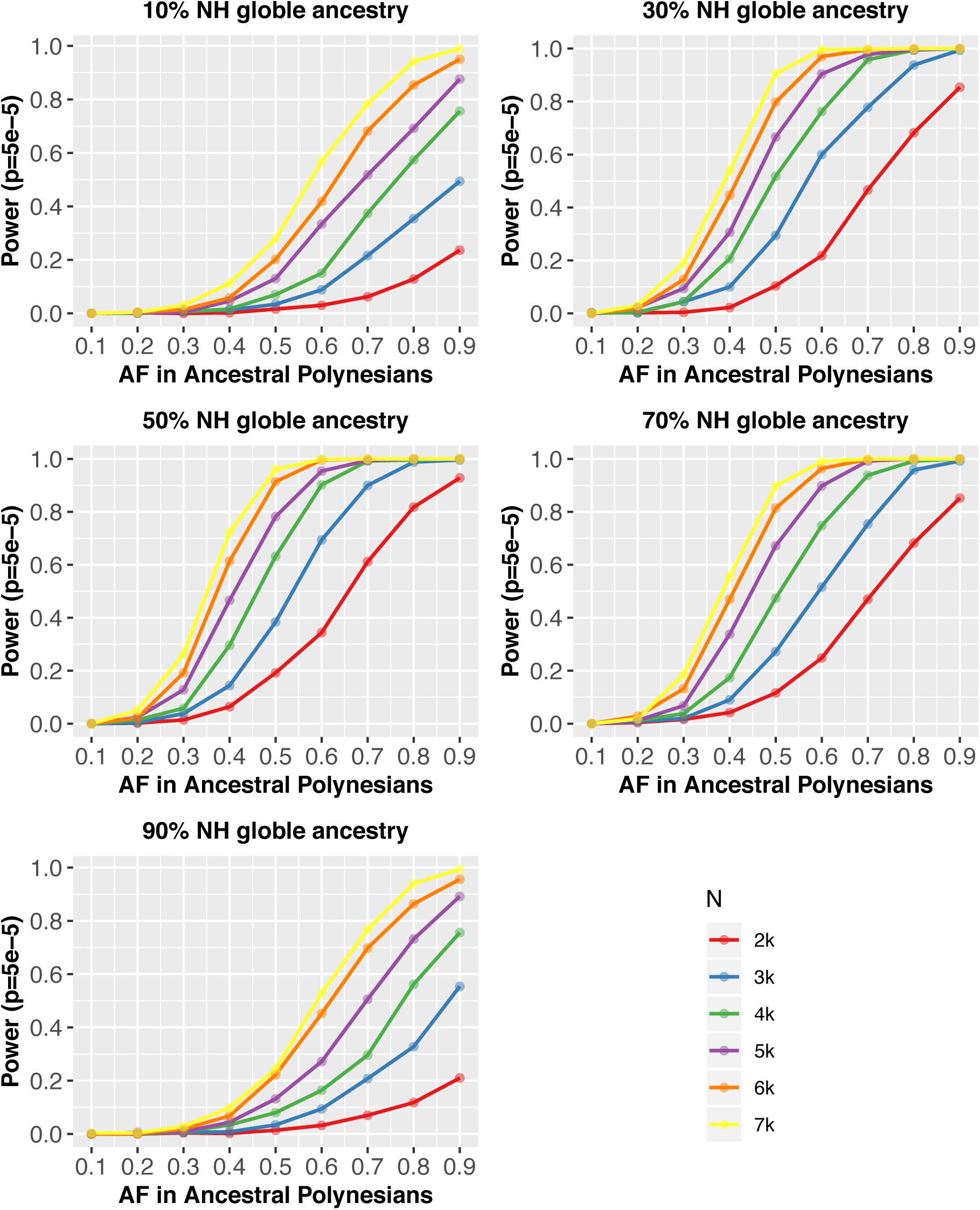
Statistical power of admixture mapping to discover *CREBRF* association with BMI. Power is estimated through simulation given a range of allele frequencies in ancestral Polynesians and sample sizes. The effect sizes and ancestry proportions were assumed using rs373863828 as example (Methods). The significance threshold for genome-wide discovery via admixture mapping is set at 5e-5.

**Figure S6.**
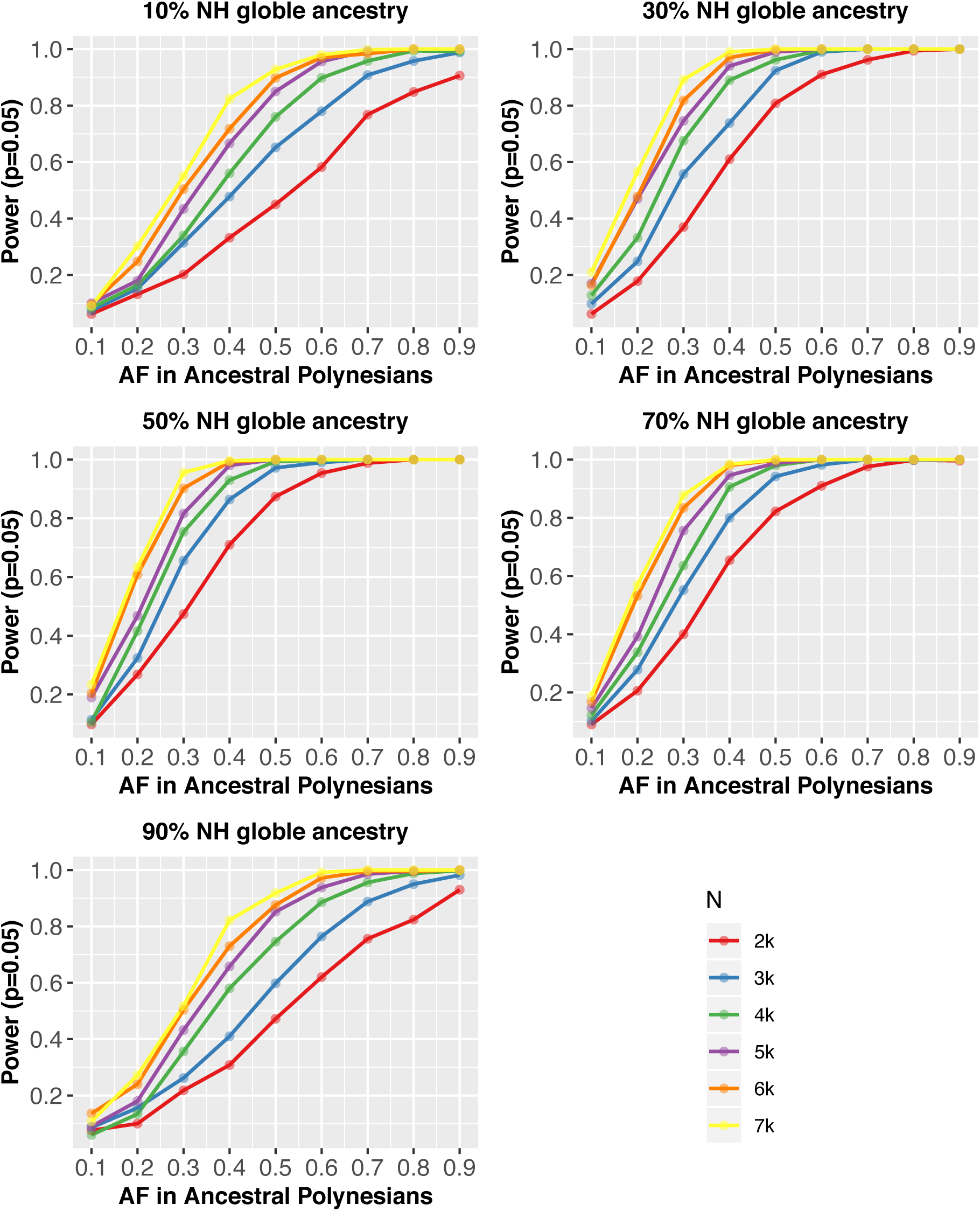
Statistical power of admixture mapping to replicate association of *CREBRF* with BMI. Power is estimated through simulation given a range of allele frequencies in ancestral Polynesians and sample sizes. The effect sizes and ancestry proportions were assumed using rs373863828 as example (Methods). The significance threshold for genome-wide discovery via admixture mapping is set at 0.05.

**Figure S7.**
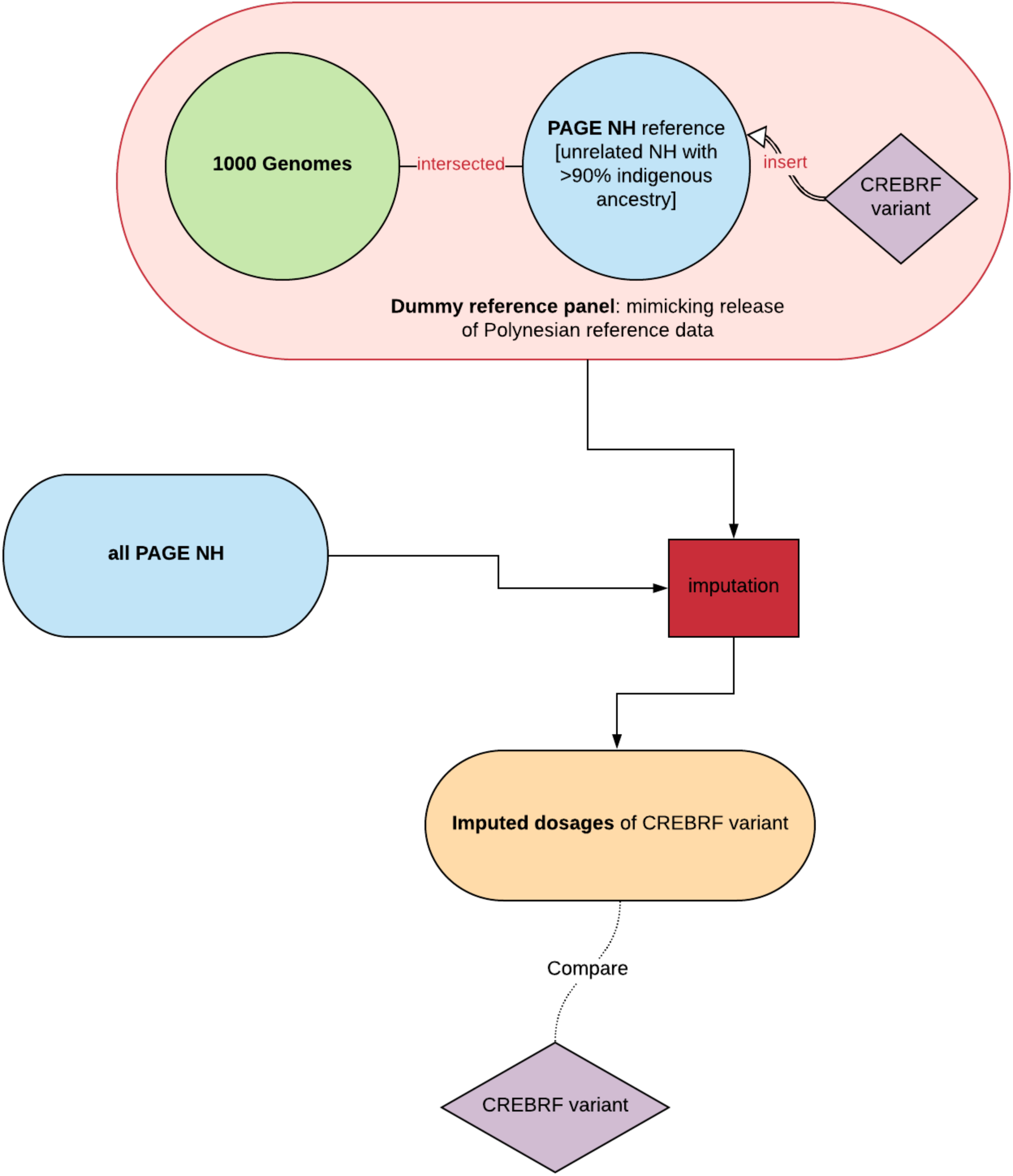
Analysis design to evaluate the imputation quality of rs373863828 using a reference panel containing internally constructed reference panel.

**Figure S8.**
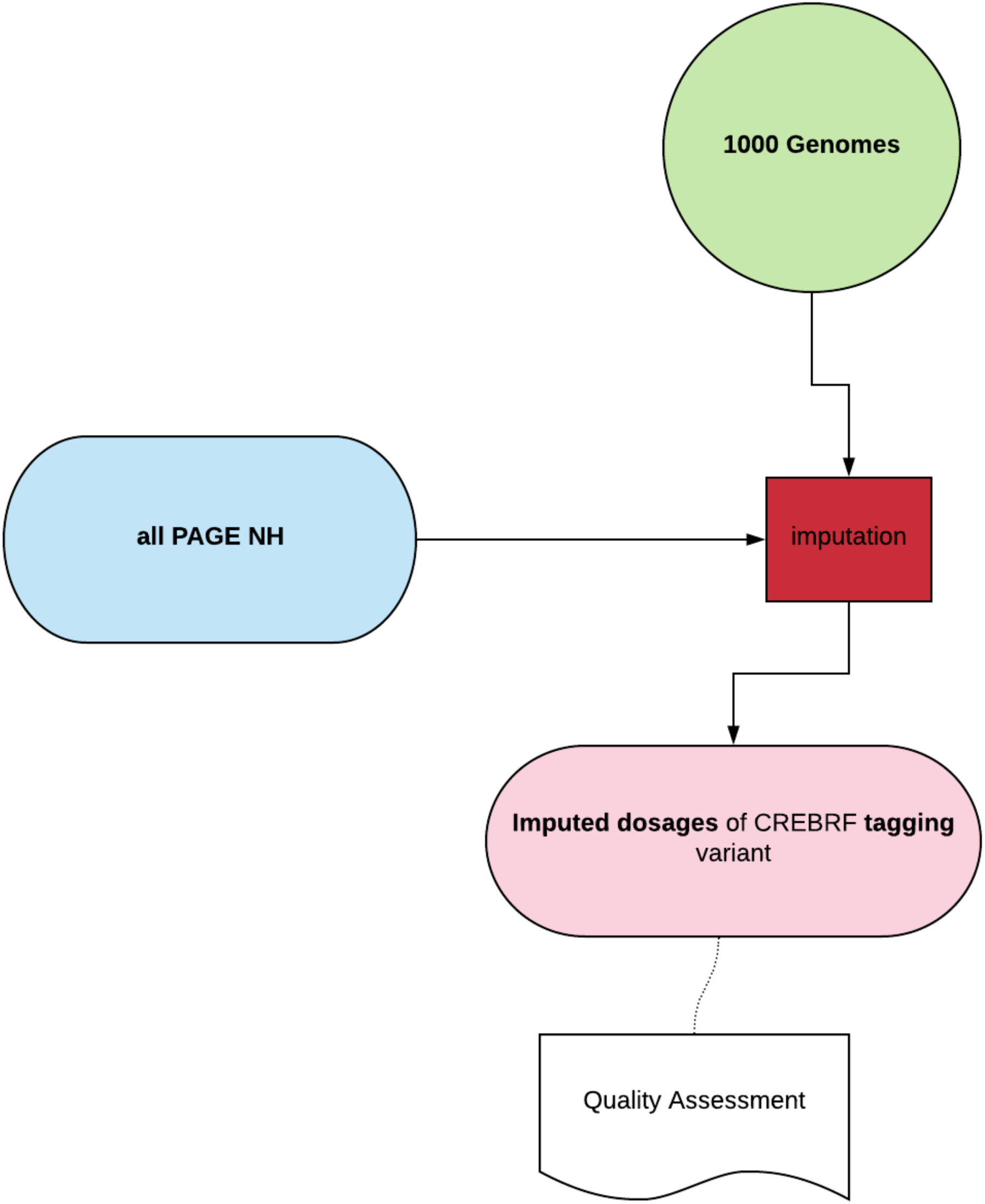
Analysis design to evaluate the imputation quality of rs12513649 using 1KGP as reference panel.

**Figure S9.**
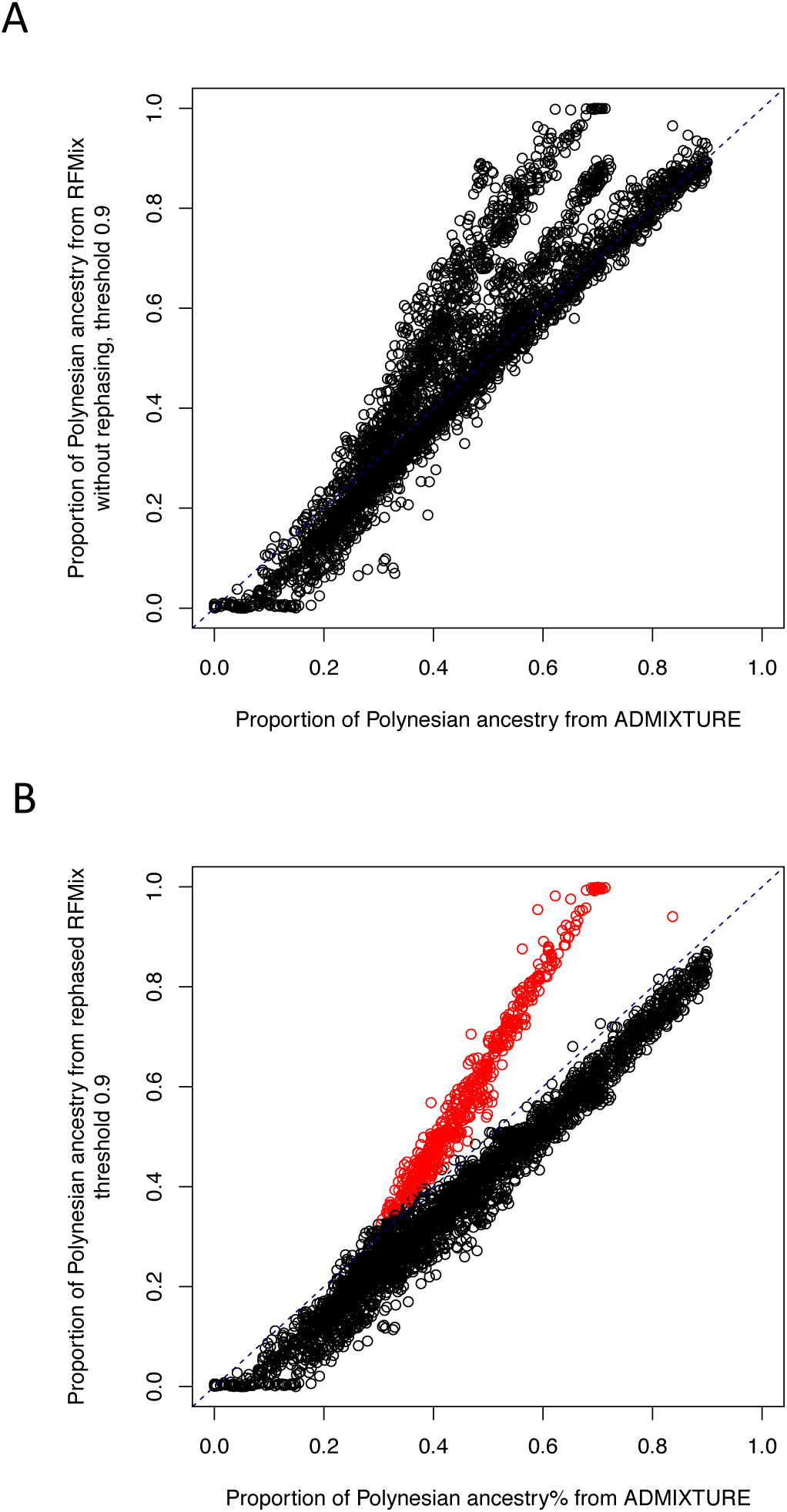
Comparison of global Polynesian ancestry estimated from ADMIXTURE and RFMix output (A) before and (B) after rephasing. RFMix calls with ancestry probability <0.9 were discarded before summing across the genome to calculate global ancestry. After rephasing, individuals with elevated RFMix estimate from that of ADMIXTURE were labeled in red; these individuals have elevated global ancestry estimate from RFMix because they are more related to the 178 reference individuals (Method).

**Table S1.** Sample size and genotype information across datasets containing Native Hawaiians

| Sample size |  |  |  |  |  |  |
| --- | --- | --- | --- | --- | --- | --- |
|  | PAGE | CRC | Smokers | NHBC | Obesity | rs373863828 |
| PAGE | 3940 | 100 | 238 | 123 | 44 | 3188 |
| CRC | 100 | 266 | 1 | 1 | 2 | 123 |
| Smokers | 238 | 1 | 318 | 14 | 0 | 281 |
| NHBC | 123 | 11 | 14 | 492 | 0 | 483 |
| Obesity | 44 | 2 | 0 | 0 | 307 | 150 |
| rs373863828 | 3188 | 123 | 281 | 483 | 150 | 3693 |

Please refer to the Method for descriptions of these datasets. The row rs373863828 reports the number of individuals from each genome-wide array dataset that had the CREBRF variant directly genotyped. PAGE dataset contained ~1,370K variants; CRC dataset contained ~256K variants; Smokers dataset contained ~269K variants; NHBC dataset contained ~152K variants; Obesity dataset contained ~1,093K variants.

**Table S2.** Phenotype transformation and covariates for linear mixed model associations.

| Traits | Phenotype Conversion | Outliers removed | Fixed covariates |
| --- | --- | --- | --- |
| BMI | sex-specific INT adjusted by age | beyond 6sd | top 10 PCs |
| Height | sex-specific INT adjusted by age | beyond 6sd | top 10 PCs |
| Waist circ | sex-specific INT adjusted by age | beyond 6sd | top 10 PCs |
| Hip circ | sex-specific INT adjusted by age | beyond 6sd | top 10 PCs |
| Waist -hip ratio | sex-specific INT adjusted by age | beyond 6sd | top 10 PCs |
| fasting glucose | INT adjusted by age at glucose draw, sex, age * sex, smoking status, bmi | t2d, no fasting, glucose < 7mmol/L | top 10 PCs |
| fasting insulin | INT adjusted by age at insulin draw, sex, age * sex, smoking status, bmi | t2d, no fasting, glucose < 7mmol/L | top 10 PCs |
| HOMA-IR | INT adjusted by age , sex, age * sex, smoking status, bmi | t2d, no fasting, glucose < 7mmol/L, diabetics | top 10 PCs |
| Adiponectin | sex-specific INT adjusted by age | beyond 6sd | top 10 PCs, log-transformed BMI |
| total cholesterol | medication adjustment as in Wojcik et al. | no fasting | age at lipid draw, sex, top 10 PCs |
| triglycerides | medication adjustment as in Wojcik et al. | no fasting | age at lipid draw, sex, top 10 PCs |
| HDL | medication adjustment as in Wojcik et al. | no fasting | age at lipid draw, sex, top 10 PCs |
| LDL | medication adjustment as in Wojcik et al. | no fasting | age at lipid draw, sex, top 10 PCs |
| Obesity | Adult > 32 kg/m2 BMI | na | age, sex, top 10 PCs |
| Diabetes | na | controls with glucose > 7 mmol/L | age at diabete, sex, bmi, top 10PCs |
| total fat mass | sex-specific INT adjusted by age | beyond 6sd | top 10 PCs |
| lean mass in legs | sex-specific INT adjusted by age | beyond 6sd | top 10 PCs, total lean mass |
| lean mass in arms | sex-specific INT adjusted by age | beyond 6sd | top 10 PCs, total lean mass |
| total lean mass | sex-specific INT adjusted by age | Beyond 6sd | top 10 PCs |
| total fat percentage | sex-specific INT adjusted by age | beyond 6sd | top 10 PCs |
| percentage of trunk fat | sex-specific INT adjusted by age | beyond 6sd | top 10 PCs, total fat percentage |
| liver fat percentage | sex-specific INT adjusted by age | beyond 6sd | top 10 PCs, total fat mass |
| visceral fat | sex-specific INT adjusted by age | beyond 6sd | top 10 PCs, total fat mass |
| subcutaneous fat | sex-specific INT adjusted by age | beyond 6sd | top 10 PCs, total fat mass |
| abdominal fat | sex-specific INT adjusted by age | beyond 6sd | top 10 PCs, total fat mass |
| HF | na | na | age, sex, top 10 PCs, (bmi) |
| Hyperlipidemia | na | na | age, sex, top 10 PCs, (bmi) |
| Hypertension | na | na | age, sex, top 10 PCs, (bmi) |
| IHD | na | na | age, sex, top 10 PCs, (bmi) |
| Stroke/TIA | na | na | age, sex, top 10 PCs, (bmi) |

For BMI, obesity, hip circumference, waist circumference, waist-hip ratio, type 2 diabetes, total cholesterol, fasting glucose, adiponectin, HDL, LDL, hypertension, and fasting insulin, we followed the guideline used in PAGE to transform the phenotypes. For HOMA-IR was calculated as $\text{HOMA-IR} = \text{fasting insulin (microU/L)} \times \text{fasting glucose (nmol/L)} / 22.5$ , and transformed per the
guideline in Minster et al<sup>1</sup>. For the remaining quantitative traits, we generally stratified by sex, adjusted by covariates such as age, and inverse normally transformed (INT) the residuals to be used in association studies.

**Table S3.** Summary of untransformed quantitative phenotypes used in association analyses

| Traits<sup>1</sup> | Mean | S.D. |
| --- | --- | --- |
| bmi (kg/m<sup>2</sup>) | 29.02 | 5.96 |
| height (cm) | 168.11 | 9.40 |
| waist (cm) | 98.59 | 15.41 |
| hip (cm) | 108.51 | 13.91 |
| waist_hip_ratio | 0.91 | 0.08 |
| glucose (mmol/L) | 4.71 | 0.68 |
| insulin (uIU/mL) | 7.85 | 6.41 |
| adiponectin (ng/mL) | 6.90 | 4.94 |
| total cholesterol (mg/dL) | 193.63 | 39.38 |
| triglycerides (mg/dL) | 122.04 | 79.72 |
| hdl (mg/dL) | 42.02 | 15.31 |
| ldl (mg/dL) | 127.62 | 36.67 |
| total fat mass (kg) | 22.64 | 7.11 |
| total lean mass (kg) | 15.51 | 3.85 |
| lean fat arms mass (kg) | 5.74 | 1.79 |
| total fat percentage (%) | 31.15 | 7.01 |
| liver fat percentage (%) | 5.80 | 4.71 |
| visceral fat area (cm<sup>2</sup>) | 166.59 | 77.71 |
| subcutaneous fat (cm<sup>2</sup>) | 235.90 | 91.27 |
| abdominal fat (cm<sup>2</sup>) | 375.79 | 132.83 |

<sup>1</sup>Outliers as defined in Table S2 are discarded.

**Table S4.** Associations of rs373863828 with adiposity-related traits from PAGE cohort

| Quantitative traits |  |  |  |  |  |  |
| --- | --- | --- | --- | --- | --- | --- |
|  | Typed genotype |  |  | Imputed dosages |  |  |
| | N | $\beta$ | P | N | $\beta$ | P |
| BMI (kg/m<sup>2</sup>) | 3185 | 0.214 | **7.55E-05** | 3936 | 0.248 | **1.50E-07** |
| Waist circumference (cm) | 2392 | 0.215 | **8.70E-04** | 3063 | 0.222 | **5.15E-05** |
| Hip circumference (cm) | 2392 | 0.206 | **1.27E-03** | 3063 | 0.254 | **3.07E-06** |
| Waist-hip ratio | 2392 | 0.00835 | 0.209 | 3063 | 0.0284 | 0.617 |
| Height | 3187 | 0.182 | **3.96E-04** | 3938 | 0.137 | **2.43E-03** |
| Fasting glucose (mmol/L) | 746 | -0.165 | 0.148 | 1146 | -0.202 | 0.0307 |
| Fasting insulin (pmol/L) | 951 | -0.0265 | 0.791 | 1557 | -0.0354 | 0.649 |
| HOMA-IR | 939 | -0.0889 | 0.378 | 1536 | -0.0778 | 0.322 |
| HDL (mg/dL) | 1230 | -0.931 | 0.493 | 1909 | -0.599 | 0.578 |
| LDL (mg/dL) | 1225 | -1.13 | 0.739 | 1897 | 0.775 | 0.772 |
| Adiponectin | 1251 | -0.122 | 0.155 | 1925 | -0.028 | 0.683 |
| Total Cholesterol (mg/dL) | 1233 | 1.48 | 0.673 | 1912 | 1.56 | 0.575 |
| Triglycerides (mg/dL) | 1233 | -5.65 | 0.428 | 1912 | -4.81 | 0.395 |
| Dichotomous traits |  |  |  |  |  |  |
|  | Typed genotype |  |  | Imputed dosages |  |  |
|  | N (case, control) | OR | P | N | OR | P |
| Obesity | 822, 2364 | 1.096 | **9.52E-05** | 955, 2983 | 1.106 | **9.72E-07** |
| Type II Diabetes | 1241, 1635 | 0.935 | 0.0162 | 1534, 2019 | 0.911 | **1.43E-04** |

Bold numbers pass Bonferroni correction of multi-trait associations: 2.5E-3

**Table S5.**
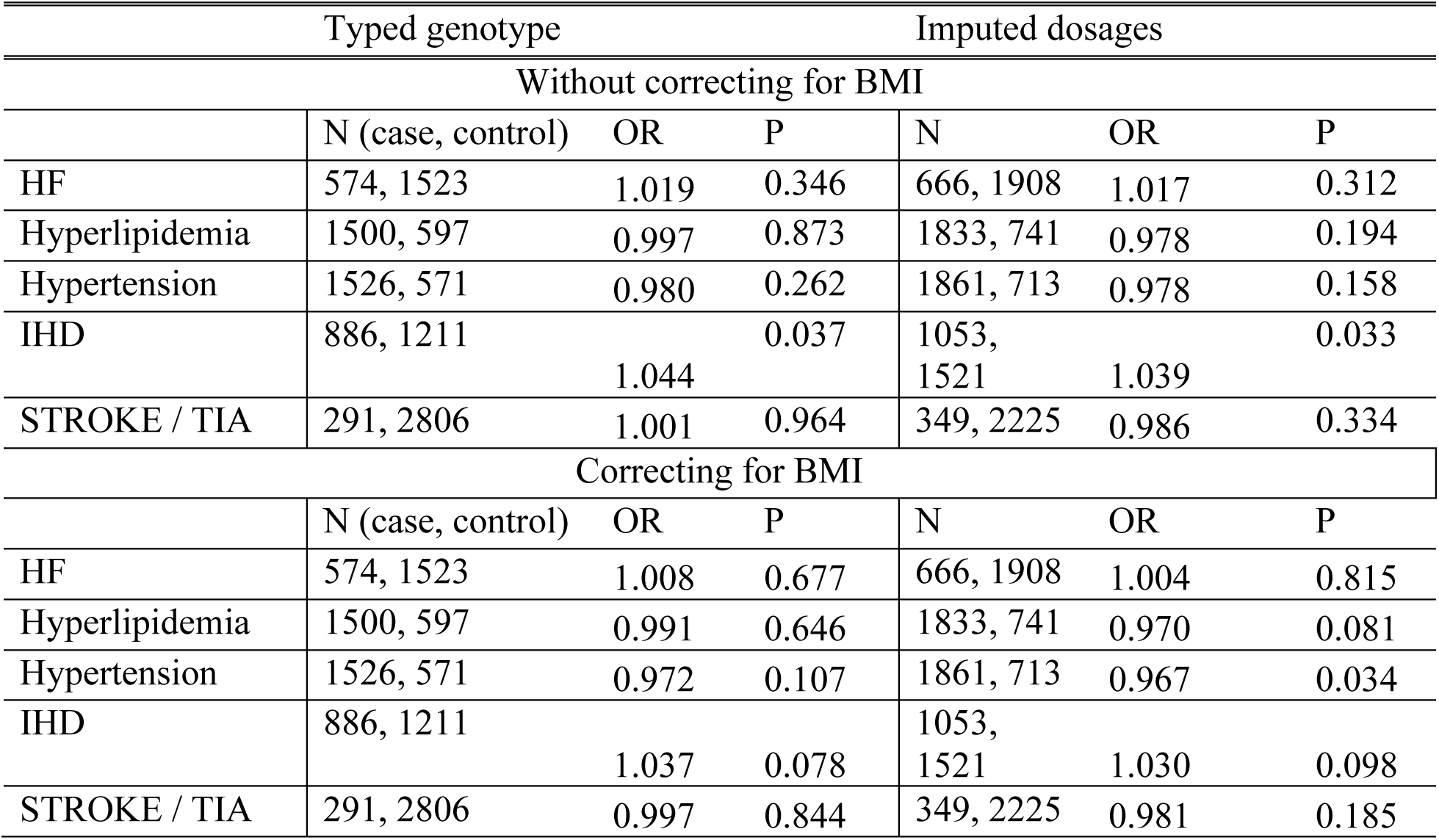
Associations of rs373863828 with cardiovascular diseases from Medicare FFS in PAGE cohort

**Table S6.** Imputation quality of rs15213649 across array platforms

|  | **PAGE** | **CRC** | **Smokers** | **NHBC** | **Obesity** |
| --- | --- | --- | --- | --- | --- |
| Overall Performance |  |  |  |  |  |
| MiniMac R<sup>2</sup> | 0.35 | 0.35 | 0.36 | 0.31 | Genotyped |
| Proportion of GP > 0.9 | 0.82 | 0.79 | 0.83 | 0.85 | Genotyped |
| Proportion of GP > 0.9, stratified by hard called genotype at rs12513649 |  |  |  |  |  |
| anc/anc | 0.84 | 0.82 | 0.85 | 0.87 | Genotyped |
| anc/der | 0.16 | 0.10 | 0.10 | 0.00 | Genotyped |
| der/der | 0 | NA | NA | NA | NA |
| Proportion of GP > 0.9, stratified by direct genotype at rs373863828 |  |  |  |  |  |
| anc/anc | 0.87 | 0.89 | 0.84 | 0.86 | Genotyped |
| anc/der | 0.49 | 0.4 | 0.69 | 0.8 | Genotyped |
| der/der | NA | NA | NA | NA | NA |

**Table S7.** Measures of imputation accuracies at rs373863828 when using 155 NH (with estimated Polynesian ancestry > 0.9) + 1KGP as reference

|  | PAGE | CRC | Smokers | NHBC | Obesity* |
| --- | --- | --- | --- | --- | --- |
| r<sup>2</sup><br>(s.e.) | 0.977<br>(6.7E-3) | 0.757<br>(0.1) | 0.913<br>(0.046) | 0.933<br>(0.036) | 1<br>(1.1E-5) |
| Minimac R2 | 0.984 | 0.96 | 0.946 | 0.939 | 0.995 |
| Concordance rate<br>(s.e.) | 0.974<br>(8.9E-4) | 0.912<br>(0.016) | 0.947<br>(0.0057) | 0.945<br>(0.0032) | 0.977<br>(7.2E-5) |

\*rs12513649 is genotyped.
Standard errors were calculated from bootstrapping (n=1000).

**Table S8.** Measures of imputation accuracies at rs373863828 when using randomly selected 152 NH + 1KGP as reference

|  | PAGE | CRC | Smokers | NHBC | Obesity* |
| --- | --- | --- | --- | --- | --- |
| r<sup>2</sup><br>(s.e.) | 0.851<br>(0.007) | 0.466<br>(0.097) | 0.899<br>(0.035) | 0.863<br>(0.044) | 1<br>(1.1E-4) |
| Minimac R2 | 0.982 | 0.962 | 0.956 | 0.964 | 0.995 |
| Concordance rate<br>(s.e.) | 0.984<br>(9.6E-4) | 0.917<br>(0.016) | 0.952<br>(0.0043) | 0.949<br>(0.0037) | 0.993<br>(1.9E-4) |

\*rs12513649 is genotyped
Standard errors were calculated from bootstrapping (n=1000).

